# High throughput instrument to screen fluorescent proteins under two-photon excitation

**DOI:** 10.1101/2020.09.04.283572

**Authors:** Rosana S. Molina, Jonathan King, Jacob Franklin, Nathan Clack, Christopher McRaven, Vasily Goncharov, Daniel Flickinger, Karel Svoboda, Mikhail Drobizhev, Thomas E. Hughes

## Abstract

Two-photon microscopy together with fluorescent proteins and fluorescent protein-based biosensors are commonly used tools in neuroscience. To enhance their experimental scope, it is important to optimize fluorescent proteins for two-photon excitation. Directed evolution of fluorescent proteins under one-photon excitation is common, but many one-photon properties do not correlate with two-photon properties. A simple system for expressing fluorescent protein mutants is *E. coli* colonies on an agar plate. The small focal volume of two-photon excitation makes creating a high throughput screen in this system a challenge for a conventional point-scanning approach. We present an instrument and accompanying software that solves this challenge by selectively scanning each colony based on a colony map captured under one-photon excitation. This instrument, called the GIZMO, can measure the two-photon excited fluorescence of 10,000 *E. coli* colonies in 7 hours. We show that the GIZMO can be used to evolve a fluorescent protein under two-photon excitation.

## 1. Introduction

The 1990s saw both the cloning of the green fluorescent protein [1] and the first application of two-photon microscopy to neuroscience [2]. Fluorescent proteins (FPs) were promising as genetically encoded probes, and two-photon microscopy made it possible to image fluorescence deep in living tissue, featuring confined excitation and decreased scattering due to the near-infrared excitation light. Today, their combination is ubiquitous in neuroscience. FPs have been engineered to sense Ca^2+^ and other indicators of brain activity [3], and two-photon microscopes have been engineered for faster imaging rates and larger fields of view to image these deep in the brains of behaving mice [4–8]. Their combination can make an even greater impact if FPs are optimized for two-photon excitation. While there have been extensive efforts to improve FP-based probes, these efforts have been almost exclusively under one-photon excitation.

Because of different molecular parameters involved in the quantum mechanical description of absorption, the two-photon properties of an FP are not equivalent to its one-photon properties. For example, the popular FP mNeonGreen [9] is two times brighter molecularly than EGFP under one photon excitation, but it is two times dimmer under two-photon excitation [10]. Often, the peak two-photon absorption wavelength is blue-shifted compared to twice the one-photon peak wavelength, due to an enhancement of a vibronic transition in the former [11]. Photobleaching rates are also variable between the two modes of excitation [12,13].

FP properties are malleable, and from day one they have been improved through directed evolution [14–16]. This involves screening a library of mutants for desired properties and creating a new library from the selected mutants. Then, the process is repeated until a significantly improved variant is found. Typically, this is done in *E. coli* colonies because they can easily express FPs, and each colony represents a single mutant. Setting up a screen under one-photon excitation is relatively simple: a colored flashlight can light up an entire plate of colonies. In contrast, two-photon excited fluorescence is confined to the focus of a pulsed, femtosecond laser, giving two-photon microscopy its edge for imaging thick samples, but creating a challenge for high throughput screening.

To date, only two efforts to evolve FPs specifically for two-photon microscopy have been described. One was a successful attempt to red shift the two-photon absorption of the green FP-based Ca^2+^ indicator jGCaMP7 [17] to better match wavelengths common to inexpensive high-powered femtosecond fiber lasers [18]. Screening of FP-expressing *E. coli* colonies was still performed under one-photon excitation with ratiometric emission registration to detect red shifted mutants. The second was a setup that could screen a plate of *E. coli* colonies under two-photon excitation by focusing the laser into a line and scanning it across the plate, while capturing the image with a camera [19]. This setup requires laser powers over 500 mW at the sample, limiting its application to expensive femtosecond amplifier systems or fixed-wavelength lasers such as fiber lasers.

Screening FPs under two-photon excitation with high throughput and adaptability is largely a hardware and software engineering challenge. We present a new instrument for this purpose called the GIZMO (gadget for implementing zippy multiphoton optimization). It is compatible with affordable, wavelength-tunable low power femtosecond lasers and can screen 10,000 *E. coli* colonies in 7 hours.

## 2. Methods

### 2.1. Experimental parameters

For the experiments described in Section 3.3, cyan to green-emitting FPs were measured. Accordingly, at the one-photon arm, a blue LED illuminating ring (455 nm, Advanced Illumination) was used with a 525/50 nm bandpass filter (Semrock) before the camera (Grasshopper3 CCD, Point Grey). At the two-photon arm, the laser (Insight DeepSee, Spectra-Physics) was set to 840 nm. The PMT (Hamamatsu) on the green side of a custom dichroic (reflects 410-550 nm, transmits 580-1000 nm, Chroma) was used with an additional 694 nm shortpass filter (Semrock). The laser power after the objective (NA 0.45, Olympus LCPLN 20XIR, working distance 8.18 - 8.93 mm) was ~5 mW. The diameter of the log spiral galvo scan pattern (see Section 3.3.1) was set at 200 μm, with only one galvo mirror active.

### 2.2. Agar plates preparation

Square, 100 x 100 mm^2^ polystyrene petri dishes were purchased from Simport (Catalog number D210-16). ~0.25 g/L of activated charcoal powder was added to LB agar (Millipore Sigma) before autoclaving, to create a non-reflective background for imaging. Each plate was filled with 80 ml of LB agar at 50° and set on a level surface overnight. This volume of agar was necessary to screen the entire plate without the objective hitting the sides of the plate. Before storing the plates, condensation was wiped off the lids with paper towels. If necessary, the inside edges of the plate were similarly dried immediately before plating. Transformed *E. coli* were spread onto the plate using 3 mm glass beads (Fisher Scientific).

### 2.3. Library preparation and analysis

To create mutant libraries, error-prone PCR was performed with the GeneMorphII kit (Agilent Technologies) or Taq polymerase (New England Biolabs) supplemented with MnCl_2_. Starting from the second round, each round of evolution also included a library created by gene-shuffling the selected clones using the staggered extension PCR method [20] and Taq polymerase (New England Biolabs). The amplified DNA fragments were inserted into the constitutive bacterial expression vector pNCS (gift from Nathan Shaner) by ligation independent cloning with In-Fusion Cloning (Clontech) or NEBuilder HiFi DNA Assembly (New England Biolabs). The cloning mixture was chemically transformed into Stellar Competent Cells (Clontech) and plated onto square agar plates for a total of ~10,000 colonies per round. The plates were incubated overnight at 37°. They were screened with the GIZMO the next day or stored at 4° for 1-4 days before screening.

The results from scanning each plate with the GIZMO were analyzed independently with a custom MATLAB program. For each colony scan, the program checks that there was a colony in the scan (i.e., not missed due to an off Z position) and then subtracts the background. It ranks the colonies by their maximum intensity value and reports the colony indices of the top five brightest colonies on the plate. The top colony was picked based on a separate MATLAB GUI that displays the one-photon plate map and marks the desired colony upon user input of the colony index. Colony PCR was performed on each colony with OneTaq (New England Biolabs). The PCR products were purified with Nucleospin (Macherey-Nagel) as per manufacturer’s instructions and sequenced with Sanger sequencing (Eurofins or Genscript).

The sequences were then analyzed with a custom MATLAB program to select the most unique mutants. These are defined by the mutants with the greatest number of missense mutations and whose mutations do not fully overlap with any other mutants. The selected mutants from each round were used as a template for the next round of evolution.

### 2.4. Instrument design and software availability

A complete CAD model of the GIZMO as well as the software and analysis programs can be found publicly at github.com/rosanamolina/gizmo-paper. The custom software runs on top of a legacy version of ScanImage (Vidrio Technologies).

## 3. Results

### 3.1. Key specifications

To design an instrument to screen fluorescent proteins under two-photon excitation, there were five key specifications to meet. First, the instrument should be set up to screen FP-expressing *E. coli* colonies. This makes it simple to recover the plasmid encoding a particular FP, and creating a larger library is as straightforward as transforming more *E. coli*. Second, the throughput of the instrument should be at least 10,000 colonies/day, corresponding to the order of a typical library size for one round of FP evolution [14]. Third, the instrument should employ a laser point-scanning system to make it compatible with wavelength-tunable low power femtosecond lasers. Fourth, the scan path needs to encompass a depth range greater than the height of an *E. coli* colony, which can reach up to 200 μm [21]. This will ensure that the entire intensity profile of a colony can be captured and relative fluorescence intensities can be accurately compared between colonies. Fifth, there needs to be automated software control with a graphical user interface (GUI). This is a critical component to maximize throughput and ensure widespread adoption in the neuroscience community.

### 3.2. Implementation

The GIZMO was designed, custom parts machined, and assembled (Figure 1). It contains a custom holder for a 100 x 100 mm^2^ square petri plate, which sits on three stages for precise XYZ movement. To efficiently scan individual colonies across the plate with two-photon excitation, the GIZMO has two detection arms connected by a stage. The first arm employs one-photon excitation to capture a widefield image of the plate, creating a map of colony positions (Figure 2A). At the second detection arm, which is essentially a two-photon microscope, this map is transformed into stage movements along the path connecting individual colonies (Figure 2B). In this way, each colony visits the focal spot of the two-photon objective to be scanned by the laser, and no time is wasted scanning the plate where no colonies are present.

**Figure 1.**
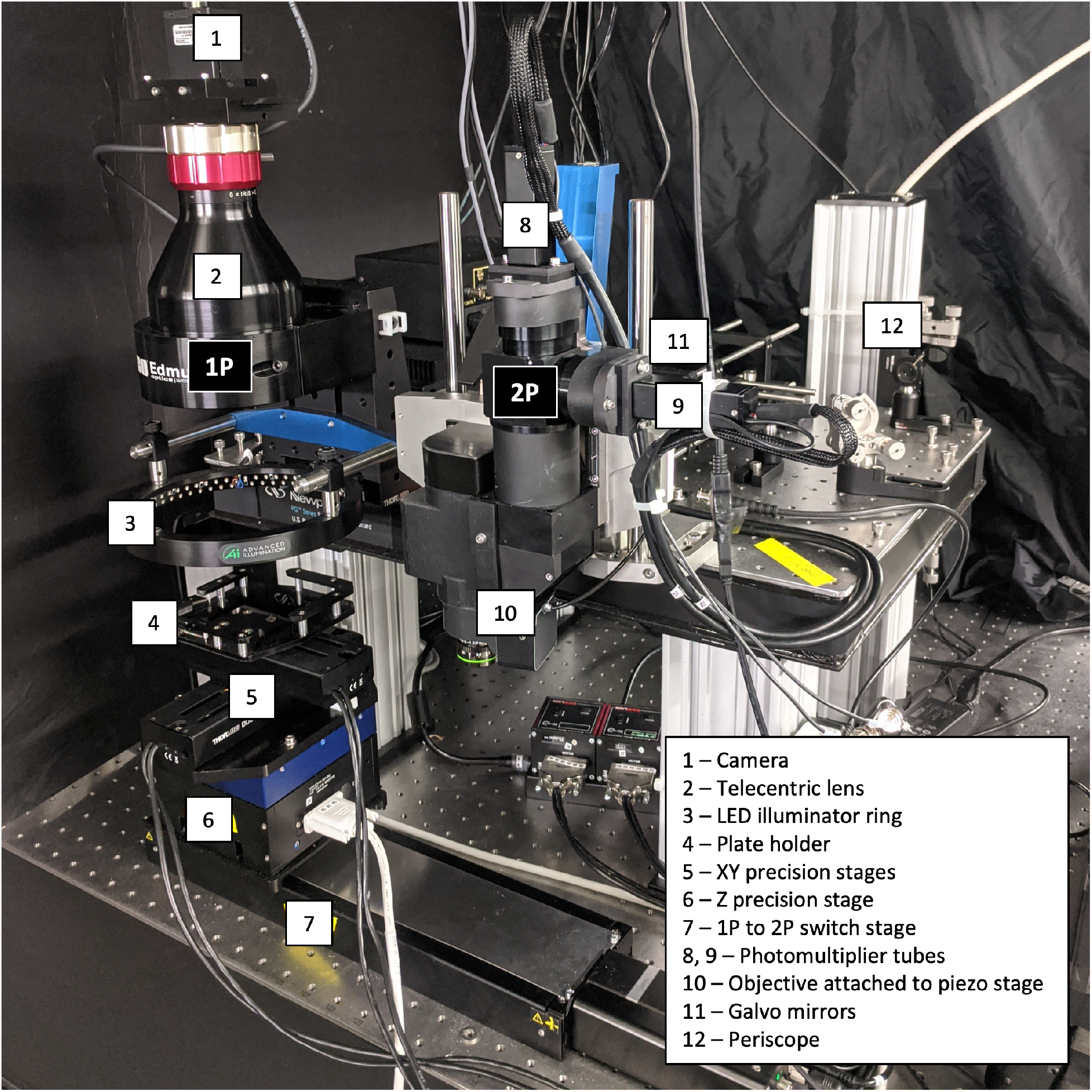
The GIZMO instrument, machined and assembled. Select components are labeled in white. The GIZMO has two detection arms (black labels): a one-photon (1P) arm creates a plate map of *E. coli* colonies (Figure 2A) to follow at the two-photon (2P) arm in order to scan each colony in an efficient manner.

**Figure 2.**
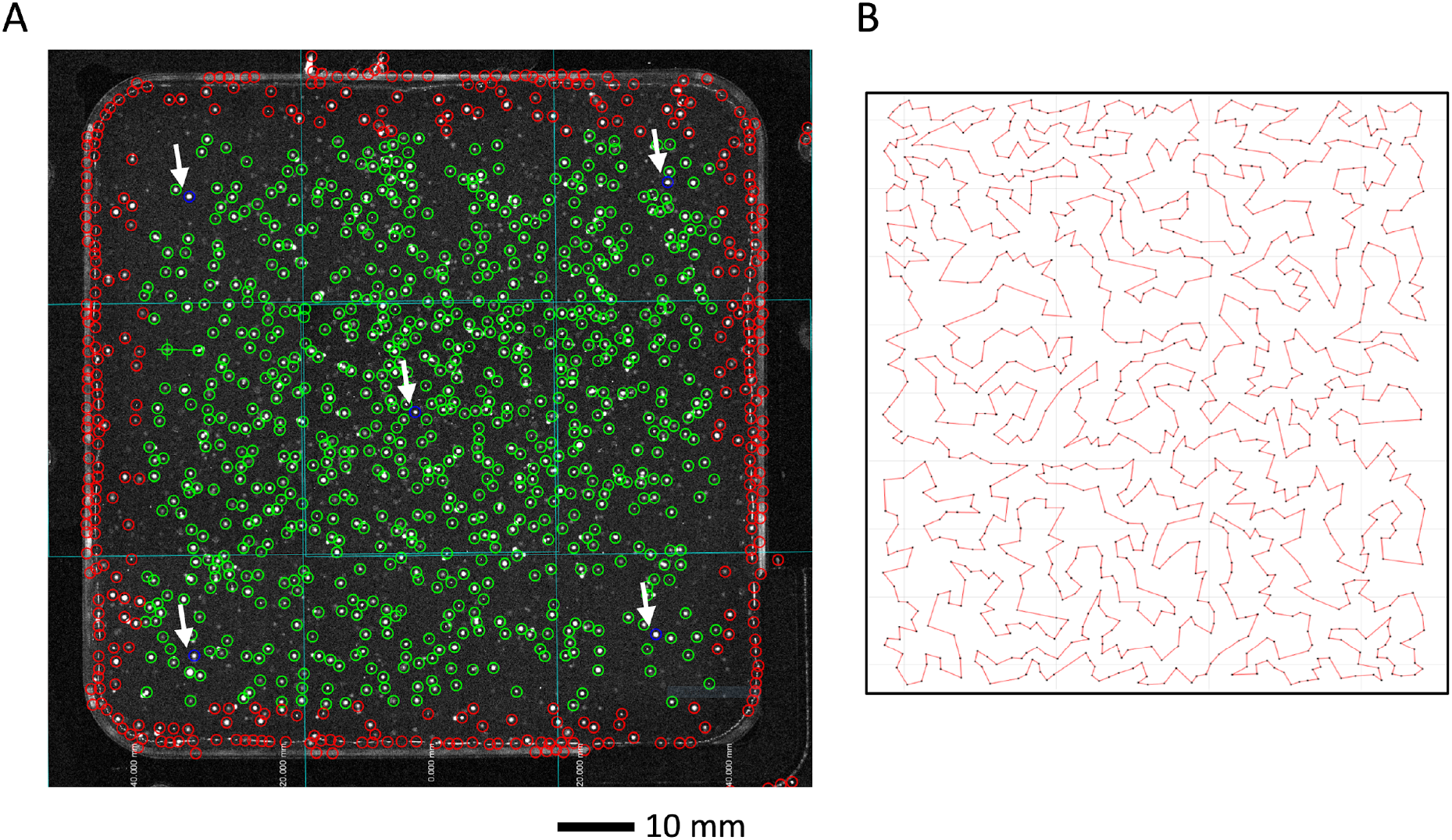
The GIZMO plate map and stage path. A) Example plate map captured by the one-photon arm of the GIZMO, as shown on the graphical user interface. The intersecting aqua lines show where the four overlapping snapshots have been automatically stitched. The open circles indicate auto-detected colonies. Green circles mark the colonies which will be scanned. The five blue circles (see white arrows) mark the test colonies which will be used to detect colony Z positions across the plate. Red circles are outside of the designated scan area and will not be scanned. B) Example of the time-efficient stage path between colonies generated from an approximate traveling salesman algorithm.

#### 3.2.1. One-photon arm

The one-photon arm captures an image of the plate under widefield excitation. The excitation light source is an LED ring illuminator (Advanced Illumination), which is exchangeable between blue (455 nm) and green (530 nm) lights. A telecentric lens (0.09X ½” GoldTL™, Edmund Optics) focuses the image through a bandpass filter onto a CCD camera (Grasshopper3, Point Grey). For a telecentric lens, the magnification of an object does not depend on its distance. Therefore, the location of each colony in the horizontal (XY) plane is accurately represented in the image despite inevitable differences in agar surface height between plates and within each plate. The field of view of the lens (71.1 x 53.3 mm^2^) does not cover the entire plate, so the software stitches together four snapshots at four XY stage positions to make the plate map (Figure 2A). Since the colonies are detected via fluorescence, those harboring nonfluorescent proteins (with mutations that caused a loss of fluorescence) will not be scanned at the two-photon arm, further saving time.

#### 3.2.2. Two-photon arm

The two-photon detection arm is set up much like a conventional two-photon laser scanning microscope. The excitation laser beam (Insight DeepSee, Spectra-Physics) first goes through a Pockels cell (Conoptics) for software-controlled laser attenuation. It is directed up to the GIZMO via a periscope and reflected off of two galvo mirrors (Cambridge Technology) for lateral scanning. It then goes through a scan lens (ThorLabs, focal length 55 mm) followed by a tube lens (Olympus U-TLUIR, focal length 180 mm). The beam transmits through a dichroic mirror (705 nm longpass, Semrock) before entering the objective (NA 0.45, Olympus LCPLN 20XIR, working distance 8.18 - 8.93 mm). The objective is mounted onto a piezo stage (Physik Instrumente) to scan the laser point through the colony quickly in Z, with a range of 400 μm. Fluorescence from the colony is collected by the objective and reflects from the dichroic to the two PMTs (Hamamatsu). The PMTs are separated by another dichroic (custom longpass, reflects 410-550 nm, transmits 580-1000 nm, Chroma) to enable two-color detection if desired. A detailed optical path diagram can be found in Figure S1.

#### 3.2.3. Software

The software is a custom implementation of ScanImage (Vidrio Technologies), written in MATLAB. In addition to controlling the hardware, it includes features critical to a timely and accurate screen of a plate. On the one-photon side, the software automatically overlays the four snapshots and detects colonies, displaying the results on a GUI (Figure 2A). There are user-adjustable thresholds for the colony-finding algorithm (e.g. maximum pixel value, minimum/maximum pixel radius of the colony) to ensure that the colonies are properly detected. A precise pixel-to-stage transform couples colony XY positions to stage movements at the two-photon side. To minimize the scan time, the software applies an approximate traveling salesman algorithm [22] to find the shortest path length between the colonies (Figure 2B).

Since the height of the agar varies between plates as well as across a single plate, colony Z positions also have to be determined. A Z-detection algorithm automatically finds the Z positions of 5 test colonies pre-selected by the user across the plate. For each test colony, the software automatically steps the Z stage in 150 μm increments and performs a 400 μm piezo scan at each step until it detects the peak of the colony. After finding the Z positions for the test colonies, the software linearly interpolates and nearest-neighbor extrapolates the Z values for the rest of the plate. With a user-determined single Z value for the entire plate, at best, only ~70% of the selected colonies lie within the 400 μm scan range of the piezo stage and are successfully captured. The Z-detection algorithm increases this to ~95%.

#### 3.2.4. Workflow and Timing

For the experienced user, scanning a total of 10,000 colonies with the GIZMO takes ~7 hours. A typical plate of 1000 fluorescent colonies takes ~3 minutes of hands-on attention (constant per plate) and ~35 minutes of automatic scanning (scaled with the number of colonies). After placing the plate into the holder at the one-photon arm, the user interacts with the GUI to take the snapshots and ensures that the autodetected colonies match what is seen by eye. The user then manually selects five test colonies for the Z-detection algorithm: one in the center of the plate, and four at the inner corners of the plate (Figure 2A). Next, the user moves the plate to the two-photon arm via the switch stage and activates the Z-detection algorithm. The final step is to scan all of the colonies, done automatically at a rate of ~2 s/colony. Stage movement between colonies takes 0.5-0.7 s/colony, and the time spent acquiring signal is 100 ms/colony. The extra time per colony present in practice is due to computational overhead, and further optimization of the software could minimize this.

The user analyzes the data with a separate MATLAB function that sorts the colonies based on the maximum intensity value and reports the colony indexes of the top five colonies. These can then be entered into a separate GUI which displays the plate and marks the colony position based on the index. The user can then pick colonies with the preferred method.

### 3.3. Calibration experiments

#### 3.3.1. Parameter considerations for colony scanning

To be able to compare the relative two-photon excited fluorescence of individual colonies, the raw scan data from each colony (exemplified in Figure 3A) is reduced to the maximum value during data analysis. To ensure that this intensity value reflects two-photon excited fluorescence, we checked the dependence of the signal intensity on the laser power. On a log-log plot of fluorescence versus laser power, the slope of a linear fit was 2.05 ± 0.03, indicating quadratic dependence and two-photon excitation (Figure S2).

**Figure 3.**
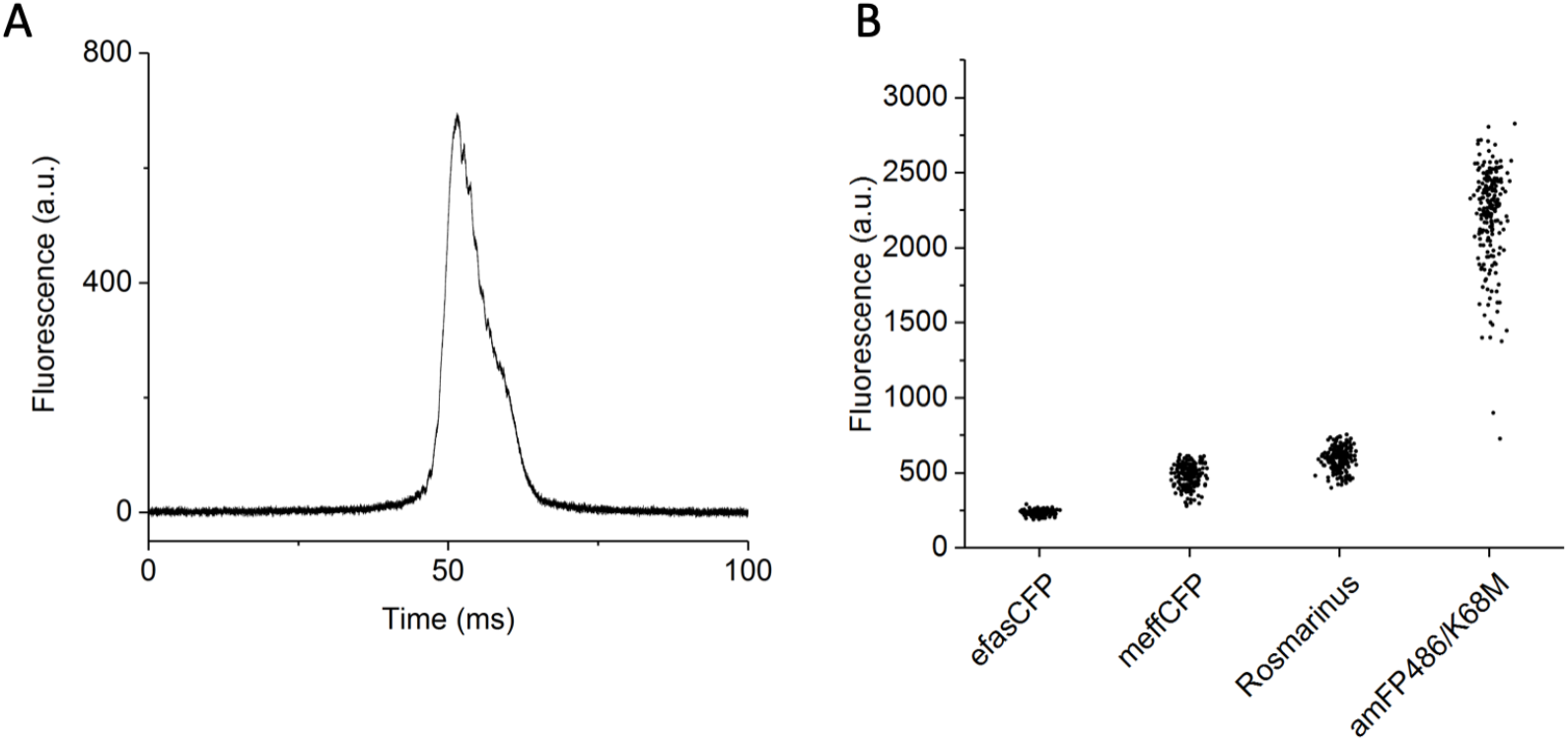
The GIZMO can distinguish between known fluorescent proteins. A) Representative raw data of a single colony scan measured by the two-photon arm of the GIZMO. The fluorescence signal detection bandwidth is ~1MHz. The relative two-photon excited fluorescence of a colony is taken as the maximum value of a scan. B) The spread of the two-photon excited fluorescence of *E. coli* colonies as measured by the GIZMO at 840 nm for four different FPs. The relative standard deviation of a single measurement for each FP (*n* colonies in parentheses) was 6.2% for efasCFP (233), 15% for meffCFP (210), 13% for Rosmarinus (195), and 14% for amFP486/K68M (210). *E. coli* were transformed with the DNA of each FP at the same time under the same conditions, and the plates were measured on the same day.

The resolution of the two-photon detection arm is not critical; we care not about the relative fluorescence of single *E. coli* bacteria (rods of diameter 1 μm and length 1-2 μm) but single *E. coli* colonies (circles of diameter 1-3 mm and height 50-200 μm). However, the smaller excitation volume of the relatively high NA objective (0.45) increases the two-photon excitation efficiency compared to a lower NA objective due to a higher power density. We measured the 840 nm wavelength beam radius at 1/e^2^ laser intensity as a function of Z with the knife edge method. Based on fitting with the equation for Gaussian beam propagation, the minimum radius of the beam was 0.91 ± 0.01 μm after the objective, corresponding to a depth of focus (Rayleigh length) of 6.2 μm (Figure S3, see Supplementary Methods).

For a single 100 ms scan of a colony, the laser point moves both axially in Z (with the piezo stage, 400 μm), and laterally in XY (with the galvo mirrors) through the colony. The galvo mirror scan pattern is set as a log spiral pattern (5 revolutions repeated 10 times) with a user-defined maximum diameter. Since a colony is radially symmetric, we found that just one galvo mirror moving is sufficient. For these measurements, lateral scanning is not as important as axial scanning. However, we found that between a diameter of 0 μm (i.e., no lateral scanning) and 200 μm, the standard deviation of the maximum intensity value across a plate of colonies expressing the same FP decreased by a factor of ~5%.

#### 3.3.2. Comparing known fluorescent proteins

Before measuring plates with libraries of mutant FPs, we determined the spread of intensity values for a plate of colonies expressing the same FP. We also asked if plates containing different FPs are distinguishable. We transformed *E. coli* with four different FPs: efasCFP, meffCFP [23], amFP486/K68M [24], and Rosmarinus [10]. A day after plating, we scanned the plates with the GIZMO (Figure 3B). The relative standard deviation of a colony measurement varied from 6.2% (efasCFP) to 15% (meffCFP). The average value for each plate with a different FP was significantly different from the others (unpaired two-sample t test, *p* < 0.001).

Two main factors contribute to the two-photon excited fluorescence intensity of a colony: the concentration of the FP and its molecular two-photon brightness. Two-photon brightness, also known as the action cross section, is the two-photon cross section (σ_2_) multiplied by the fluorescence quantum yield (φ), and it is measured in Goeppert-Mayer units (GM, 10^−50^ cm^4^ s molecules^−1^ photons^−1^). Based on the known molecular brightness of each FP at the excitation wavelength (840 nm), Rosmarinus should be the brightest of the four FPs (85 GM), then amFP486/K68M (70 GM), and then meffCFP and efasCFP (55 GM) [10]. As measured on the GIZMO, the average colony intensities of amFP486/K68M, meffCFP, and efasCFP were 3.7, 0.8, and 0.4 times that of Rosmarinus, respectively (Figure 3B). This shows that the concentration factor plays a large role in the overall difference between FPs.

#### 3.3.3. Directed evolution of Rosmarinus

Our final goal was to demonstrate that the GIZMO could be used to evolve a FP under two-photon excitation. We decided to start with Rosmarinus, as it was already molecularly two-photon bright but had room to improve in terms of functional protein concentration, as seen in the previous experiment (Figure 3B). For the screening laser wavelength, we chose 840 nm, which is slightly blue-shifted from the two-photon absorption peak of Rosmarinus (~850 nm). We based this choice on the evidence that two-photon brighter green FPs tend to be blue-shifted [10]. We created a library of Rosmarinus variants through random mutagenesis (see Methods). For the first library, we screened about 6,000 mutants and picked the top 1% brightest colonies from each plate as a template for the next round of evolution. For subsequent rounds, we picked the top colony from each plate, sequenced them, and selected 5-10 mutants based on sequence diversity. We also performed gene-shuffling to find beneficial mutation combinations (see Methods).

The two-photon excited fluorescence of the selected mutants as measured by the GIZMO steadily increased for seven rounds of evolution before leveling off at the eighth round (Figure 4A, see Figure S3 for log scale). Clone 7.02, the top clone from the seventh round, was on average 9 times brighter than Rosmarinus (Figure 4A). It contains 11 amino acid mutations relative to Rosmarinus: H13R, M24V, I40V, G128D, V131I, T136S, E182G, N212T, A219T, K225E, and K227E (numbered according to Genbank KY931461). We characterized the two-photon absorption properties of Clone 7.02 in purified protein (see Methods) and compared them to Rosmarinus. The two-photon molecular brightness and spectral shape were identical (Figure 4B). Evolving Rosmarinus under two-photon excitation with the GIZMO at 840 nm led to a dramatic increase in functional protein concentration in *E. coli* colonies but did not alter the molecular brightness.

**Figure 4.**
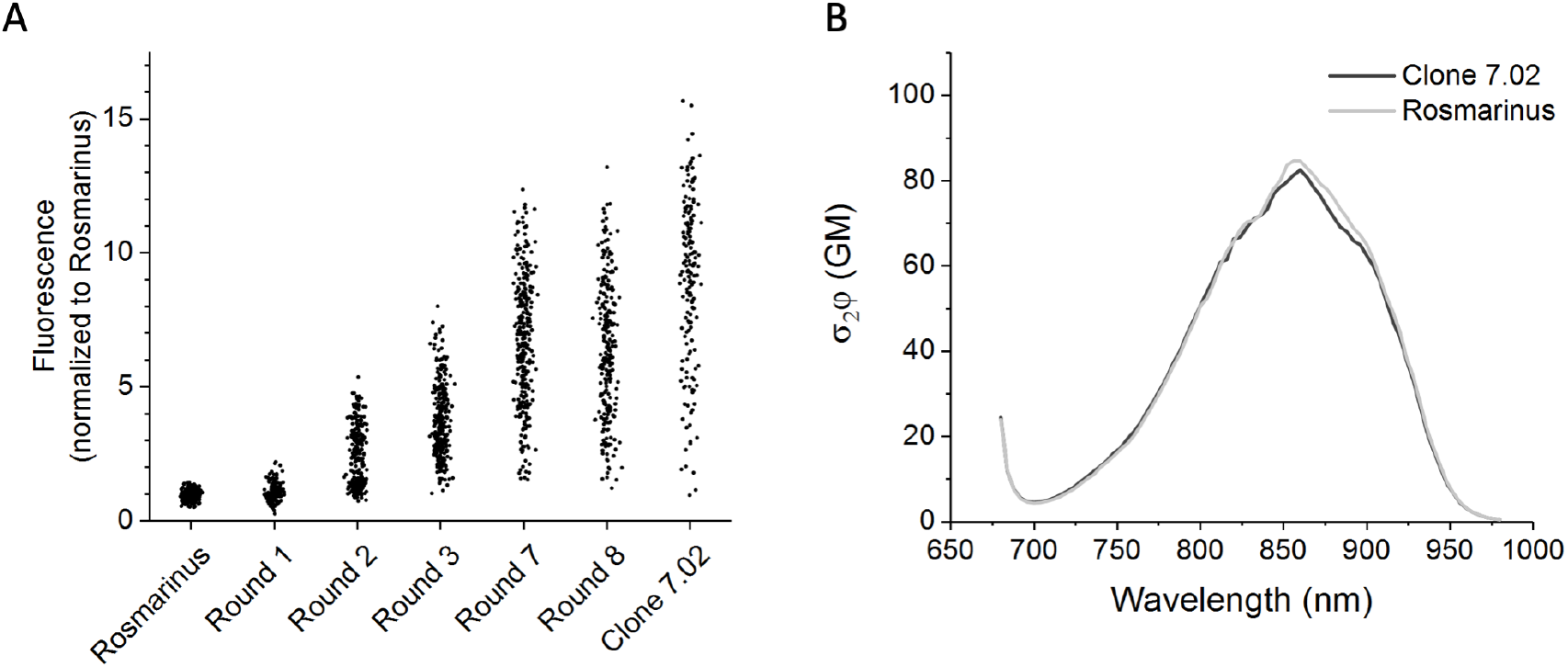
Directed evolution under two-photon excitation. A) Two-photon excited fluorescence of colonies from different rounds of evolution, excited at 840 nm. Shown are Rosmarinus, the selected mutants from each round of evolution, and the brightest mutant from Round 7 (Clone 7.02). Rounds 4, 5, and 6 are omitted for simplicity. The same data is plotted in log scale in Figure S3. B) The two-photon molecular brightness of Clone 7.02 is identical to the parent protein Rosmarinus, as measured in purified protein.

## 4. Discussion

We have designed and implemented an instrument that can measure the two-photon excited fluorescence of 10,000 *E. coli* colonies expressing a library of FPs in 7 hours. It meets the challenge of quickly scanning a laser across a plate of colonies by first taking a widefield one-photon image of the plate, which then serves as a map of colony positions to visit with the two-photon point scanning approach. The one-photon and two-photon detection arms follow standard optical design. The custom software seamlessly integrates the two arms to make fast FP evolution possible.

To test the GIZMO in action, we performed a proof-of-principle experiment to evolve Rosmarinus, a promising FP for two-photon excitation because of its strong two-photon brightness. Screening under two-photon excitation put selective pressure on the mutants that expressed better in *E. coli* and maintained a high molecular two-photon brightness, which may have been lost if the screening was done under one-photon excitation. The colonies did not get brighter after the seventh round of directed evolution, suggesting that a saturation point had been reached in terms of expression. To find mutations that increase the molecular brightness as well, it is possible that a larger library is necessary. Considering the throughput of the instrument, 50,000 mutants may still be practical. Alternatively, libraries that target specific sites in the protein predicted to be near the chromophore may be a more efficient way to find molecularly brighter mutants.

A two-photon optimal FP should be bright and express well, but it also needs to be photostable under two-photon excitation. Software development is in progress to add photobleaching characterization and screening capabilities to the GIZMO. No extra hardware is required. The photobleaching rate under two-photon excitation usually depends stronger than quadratically on the laser power [12], and it can also depend on the excitation wavelength [25,26]. These are parameters we are taking into consideration to develop an accurate and informative two-photon photobleaching screen.

In addition to photobleaching, the GIZMO has more capabilities than demonstrated here, both currently available and possible with further development. While we worked with cyan to green FPs, it is readily adaptable for other colors, including orange and red. Its compatibility with a wavelength-tunable laser makes it possible to screen for desired spectral properties. There are two PMTs for the option of including a second FP in the screen. In combination with functional assays, the GIZMO could be used to screen FP-based biosensors that are able to be expressed in *E. coli*, such as kinase sensors [27] or Ca^2+^ sensors [17,28–30]. Since it uses a pulsed excitation source, it could be fitted with a digitizer to enable fluorescence lifetime screening. It could also be adapted to screen fluorescent proteins under three-photon excitation, which can penetrate deeper into the brain than two-photon excitation [31]. The properties of the GIZMO make it a versatile instrument.

With the help of the GIZMO, protein engineers can create a new generation of genetically encoded fluorescent tools specifically optimized for two-photon microscopy. These tools will expand the accessible landscape of experiments to investigate the brain.

## Acknowledgments

We thank Jon Arnold at the Janelia Experimental Technology Department for his contributions to the mechanical design of the GIZMO; Doug Kim, Loren Looger, and Vivek Jayaraman for supporting this project at Janelia; Kaspar Podgorski and Abhi Aggarwal for keeping the GIZMO employed at Janelia; and William Pardis for setting up the GIZMO in Bozeman.

