## Supplemental Information for "High throughput instrument to screen fluorescent proteins under two-photon excitation"

### 1. Methods

#### *1.1. Beam waist measurement*

To measure the waist of the beam, a razor blade was fastened to the plate holder on the GIZMO and a power meter was centered below the objective. At various heights set by the Z stage, the power attenuation of the 840 nm laser was recorded as a function of stepping the X stage to move the razor blade across the beam. At each Z height, the beam radius $w$ at $1/e^2$ laser intensity was found by fitting the data with the following equation:

$$P(x) = P_0 + \frac{P_{max}}{2}\left(1 - erf\left(\frac{\sqrt{2}(x-x_0)}{w}\right)\right) \quad \text{(S1)}$$

Here, $P$ is the laser power measured at the X stage position $x$, $P_0$ is the background power, $P_{max}$ is the maximum power, $x_0$ is the position of the center of the beam, and *erf* is the standard error function.

The beam waist $w_0$ was found by fitting the beam radius $w$ as a function of the Z stage position $z$ with the Gaussian beam propagation equation (Figure S3):

$$w(z) = w_0\left(1 + \left(\frac{(z-z_0)\lambda}{\pi w_0^2}\right)\right)^{\frac{1}{2}} \quad \text{(S2)}$$

where $z_0$ is the position of the beam waist and $\lambda$ is the laser wavelength.

#### *1.2. Protein purification*

Proteins were purified using His60 Ni Superflow Resin (Clontech). Photophysical measurements were made in the elution buffer (50 mM sodium phosphate, 300 mM sodium chloride, 300 mM imidazole; pH 7.4).

#### *1.3. Two-photon characterization*

Two-photon characterization was done as in previous work [1,2], with a modification to the method for measuring the two-photon cross section. A femtosecond tunable InSight DeepSee laser (Spectra-Physics) was directed into a PC1 Spectrofluorimeter (ISS) containing the sample in a 3 mm cuvette. The laser was controlled with a custom LabView program to measure the spectral shape relative to the reference standard Coumarin 540A (Exciton) in DMSO (MilliporeSigma) (the spectral shape of Coumarin 540A from 680 nm to 760 nm was corrected with Prodan in DMSO [3]). A detailed description of this setup can be found in [2].

The two-photon cross sections ($\sigma_2$) were measured at excitation wavelengths of 840 nm and 900 nm, relative to fluorescein (MilliporeSigma) in 1 mM NaOH (extinction coefficient (491 nm) = 92,000 $M^{-1}cm^{-1}$; $\sigma_2$(840 nm) = 12.9 GM; $\sigma_2$(900 nm) = 15.4 GM [4]). To calculate the absolute two-photon cross sections, fluorescence signal as a function of laser power was collected under both two-photon excitation and one-photon excitation conditions. One-photon excitation was done using the 458 nm line of an argon ion laser (IMA101040 ALS, Melles Griot), selected with an interference filter. The fluorescence was collected through a 520 longpass filter (Chroma) and a 770 shortpass filter (Semrock). Collection conditions were identical between excitation conditions. The fluorescence signals as a function of laser power were fit to a parabola or a line for two-photon or one-photon excitation, respectively. The resulting fit coefficients represent a multiplication of 1) relative absorption strength at the excitation wavelength, 2) relative concentration, 3) PMT spectral sensitivity parameters, 4) laser excitation parameters, and 5) fluorescence collection efficiency. Factors 2 and 3 are identical under both excitation conditions for each sample, and factors 4 and 5 are identical for every sample. Thus, the known extinction coefficients of the samples at 458 nm and the known $\sigma_2$ of fluorescein at the two-photon excitation wavelength were used to calculate the $\sigma_2$ of the FP.

##### *1.4. Fluorescence quantum yield and extinction coefficients*

The fluorescence quantum yields were measured with an integrating sphere spectrophotometer (Quantaurus-QY, Hamamatsu). The reference (buffer-only) measurement was done in the same cuvette as the sample measurement. The extinction coefficients were measured by scanning the one-photon absorption spectra with the Lambda 950 Spectrophotometer (Perkin-Elmer) during stepwise alkaline denaturation with 1 M NaOH. The extinction coefficient of the protein was calculated relative to the known extinction coefficient of the fully denatured form of the chromophore, 44,100 $M^{-1}cm^{-1}$ [5].

#### 2. Figures

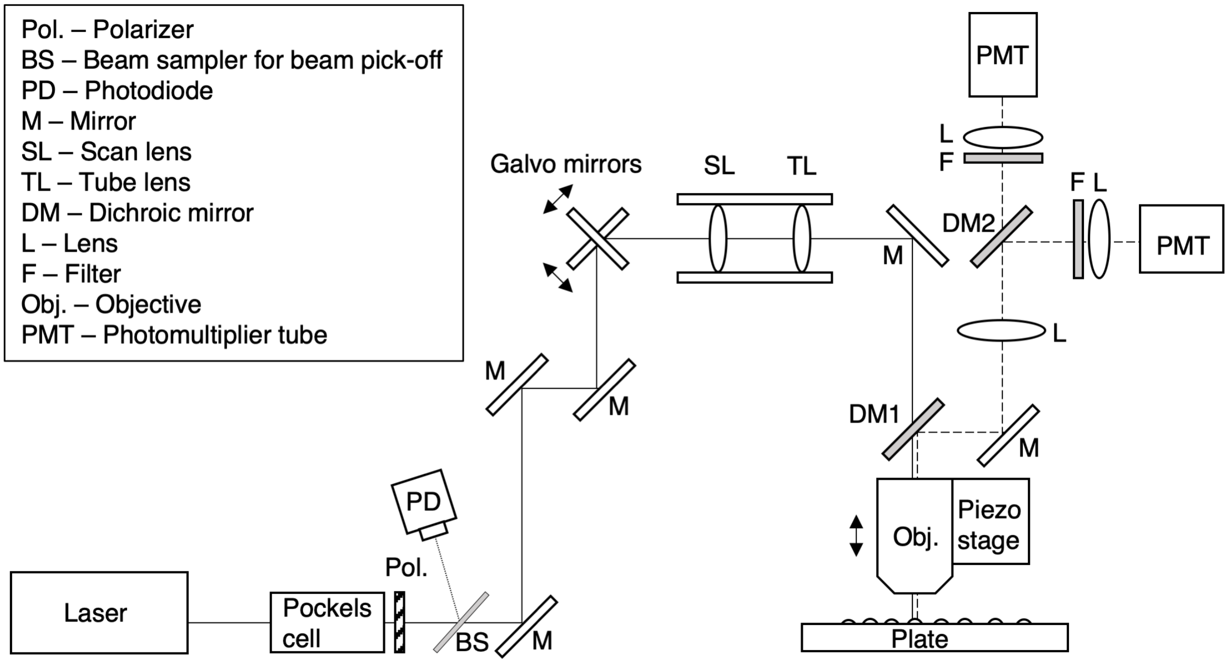


Figure S1. **Optical path of the two-photon detection arm.** The excitation path of the laser is shown with a solid line, and the emission path of the fluorescence is shown with a dashed line.

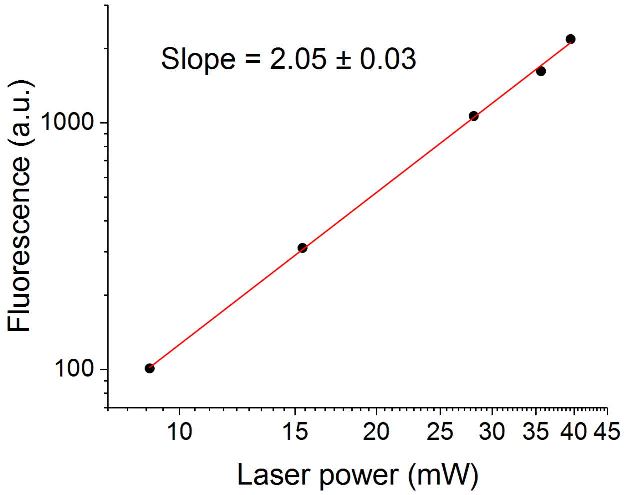


Figure S2. **Quadratic power dependence of fluorescence signal.** The maximum fluorescence signal measured by the GIZMO of an *E. coli* colony expressing Rosmarinus was quadratically dependent on the laser power.

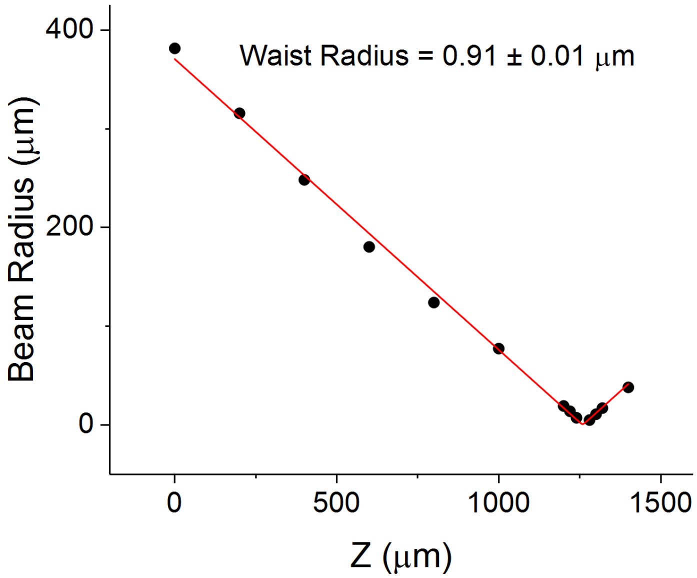


Figure S3. **Beam waist measurement**. The beam radius at $1/e^2$ laser intensity was determined at various Z heights with the knife edge method. The red line shows the Gaussian beam propagation equation fit for the laser wavelength 840 nm (Equation S2).

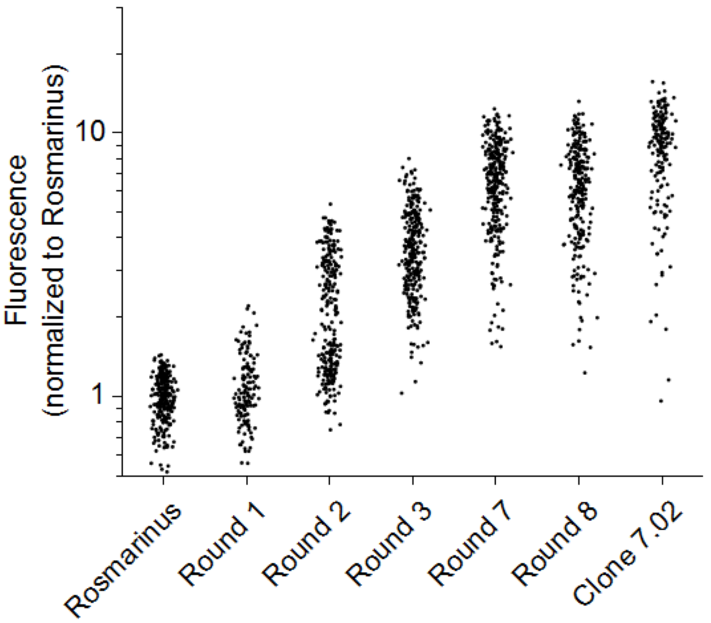


Figure S4. **Directed evolution experiment to evolve Rosmarinus**. The data here are the same as Figure 4a of the main text but plotted in log scale. Shown are plates measured with the GIZMO at 840 nm of colonies expressing Rosmarinus, the selected mutants from each round of evolution, and the brightest mutant from Round 7 (Clone 7.02). Rounds 4, 5, and 6 are omitted for simplicity. The plates were transformed under the same conditions and measured on the same day.
